# The CSF transcriptome in pneumococcal meningitis reveals compartmentalised host inflammatory responses associated with mortality

**DOI:** 10.1101/2024.05.24.592913

**Authors:** José Afonso Guerra-Assunção, Probir Chakravarty, Gabriele Pollara, Cristina Venturini, Veronica S Mlozowa, Brigitte Denis, Mulinda Nyirenda, David G Lalloo, Mahdad Noursadeghi, Jeremy S Brown, Robert S Heyderman, Emma C Wall

## Abstract

**Background:** Pneumococcal meningitis (PM) has persistently poor clinical outcomes, especially in sub- Saharan Africa. To better characterise the inflammatory response and identify factors associated with mortality we compared paired peripheral blood and cerebrospinal fluid (CSF) transcriptomes before the initiation of antibiotics in Malawian adults with proven PM.

**Results:** Blood transcriptional profiles were obtained in 28 patients with PM, with simultaneous paired with CSF profiles available for 13 patients. 15/28 (52%) patients died. Comparison of the transcriptome between CSF and blood compartments showed upregulation of 2293 differentially expressed genes in CSF and 909 in blood; enriched pathways in CSF included inflammasome activity and neutrophil migration/activation in the CSF, contrasting with enrichment for pathways including platelet and endothelial activation, cell cycle, cytokine release and oxidative stress in the blood transcriptome. Comparison of CSF profiles between survivors and non-survivors revealed 1829 differentially expressed genes, non- survivor CSF was enriched for multiple innate inflammatory pathways, including IL-17A and Type 1 interferons and proteolysis. In contrast, minimal transcriptomic differences between outcome groups were detected in blood.

**Conclusion:** Inflammation in PM is characterised by compartmentalised responses in blood and CSF. Poorer outcomes are associated with an dysregulated innate immune host response to *S. pneumoniae* in the CSF compartment.

## Introduction

Despite effective conjugate vaccines and better critical care available in many settings, *Streptococcus pneumoniae* remains the most prevalent cause of community acquired bacterial meningitis globally^1–3^. The greatest burden of pneumococcal meningitis (PM) in adults and adolescents falls on low and middle income countries (LMICs) with high HIV prevalence,^4^ where the reported mortality is 50-70%^5,6^.

Animal model and human post-mortem studies indicate that the pathogenesis of PM is characterised by a marked neutrophil-mediated inflammatory response to bacterial invasion in the CSF.^7,8^ This inflammatory cascade combined with the cytotoxic effects of host pro- inflammatory mediators^9,10^ and bacterial toxins drive tissue damage characterised by apoptotic neuronal cell injury, raised intracranial pressure (ICP), thrombosis, cerebral oedema, and cerebral ischaemia^11,12^. Both higher bacterial loads and ineffective bacterial clearance by neutrophil extracellular traps (NETS) have been implicated in pathogenic inflammation in PM^13–17^, but the modifiable upstream drivers of lethal brain damage remain poorly understood.

Evidence of a prominent role for host-mediated inflammation in meningitis-associated brain damage supported the trials of anti-inflammatory agents, such as dexamethasone as adjunct therapies for PM. Dexamethasone has demonstrated efficacy in controlled trials in industrialised countries in HIV-negative adults with PM; estimated relative risk reduction in mortality of 0.5 (95% CI 0.3 – 0.83)^18^. However, all adjunctive agents including glucocorticoids have been proven to be ineffective or even harmful in patients with PM in LMICs in controlled trials (irrespective of HIV-1 serostatus)^1,19,20^. The differences in overall case fatality rates by geographical location and response to dexamethasone in patients with PM remain unexplained. Patients with bacterial meningitis in LMICs are younger, have a higher incidence of HIV co-infection, present later, have very high bacterial loads in CSF and important differences in bedside predictors of poor outcome compared to patients in better esourced settings^15,21,22^. Data from animal models and children with PM suggest excessive inflammatory responses determine disease severity,^8,11,23^ however the mechanisms of tissue damage from the host-pathogen interaction in PM remain poorly understood, and few tractable therapeutic targets have been identified to date^6,23–26^.

Transcriptional profiling in infectious diseases offers an opportunity to describe the immunomodulatory processes associated invasive infection in greater depth^27–29^. Meningitis- specific transcriptomic data in children with tuberculous meningitis (TBM) have shown compartmentalisation of neurological damage within the CNS, compartment compared to inflammasome activation in serum, whereas in neonates with bacterial meningitis (BM) caused by a range of pathogens, transcriptome data suggest activation of TLR-signalling pathways in CSF and an evolution of the cellular response over time^30,31^. Whilst informative, these reports are not directly relevant to adult PM, particularly towards amelioration of intractably poor clinical outcomes. To better understand the excessively high mortality in patients with PM in LMICs, we compared blood and CSF host transcriptomic responses in Malawian adults with proven PM on admission to hospital, testing if either the cellular composition of CSF or transcriptional response in paired blood and CSF differed between survivors and non survivors.

## Results

### PM patients

We extracted RNA of sequencing quality (*RIN* >7) from the blood of twenty-eight adults with proven PM along with paired CSF for thirteen from pre-antibiotic samples collected on admission to hospital (Figure 1). The median age of the patients was 33 (range 26-66) years, 21/25 (84%) were HIV co-infected, mortality at 6 weeks post-discharge was 15/28 (52%) (Table 1). All patients received parenteral ceftriaxone within 3 hours of arrival in hospital.^1^ Non-survivors had lower Glasgow Coma Scores on admission to hospital compared to survivors (11/15 (6-13) vs 14/15 (13-15), OR 0·13 CI 0.22 - 0.8, p=0.02), but with no differences in reported time between symptom onset to presentation. Median CSF WCC were 30 cells/mm^3^ (range 0-635), with no differences in absolute CSF WCC between survivors and non-survivors (OR 0.95, 95% CI 0.4-2.2 p=0.91) (Table 1). In contrast, CSF bacterial loads were approximately 100-fold higher in non-survivors (4.37x10^7^ DNA copies/ml, IQR 8.43x10^6^– 2.18x10^8^) than in survivors (5.7x10^5^ copies/ml, IQR 3.2x10^4^ – 4.7x10^6^) p=0.02 (Table 1).

**Figure 1:**
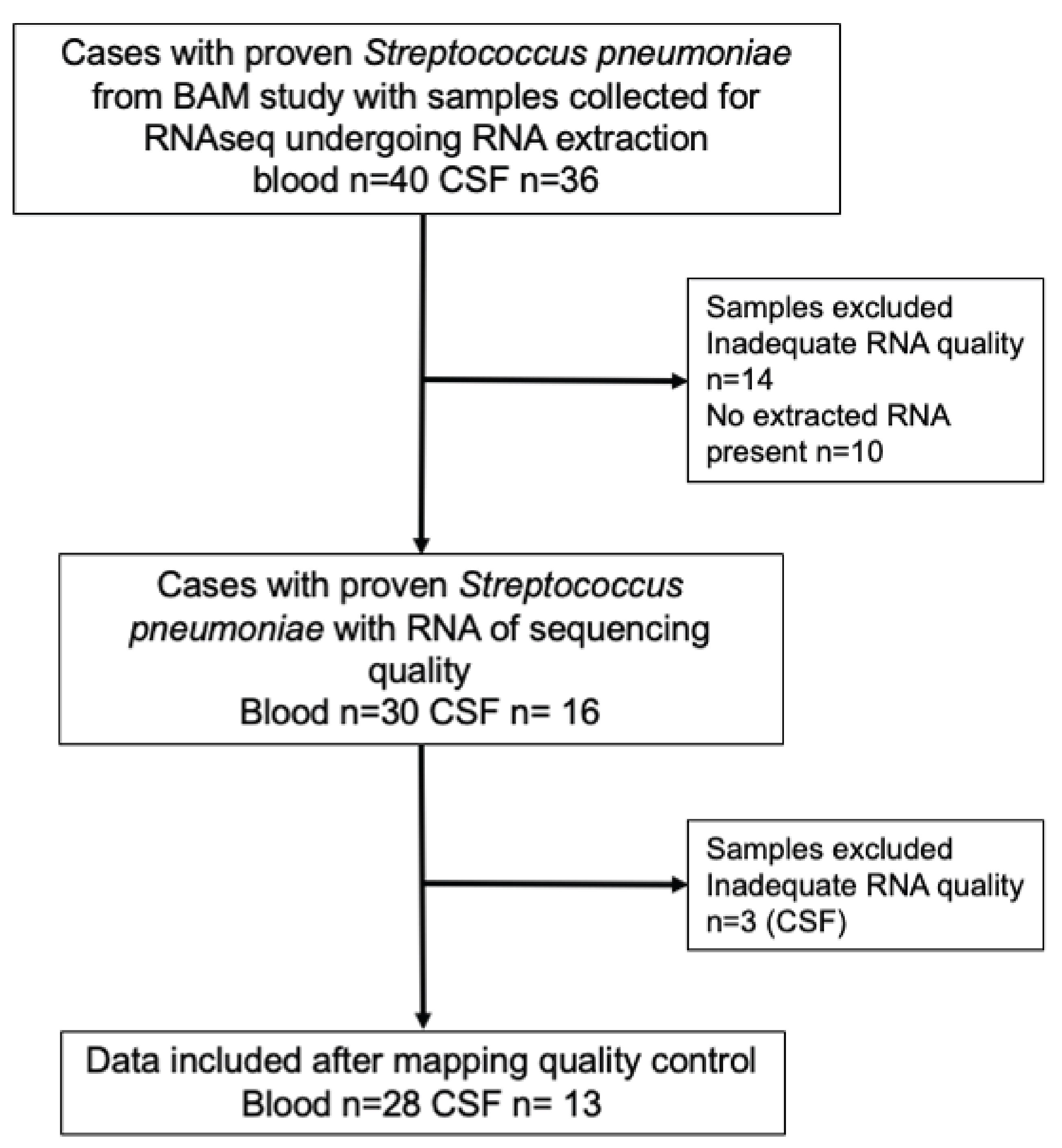
Selection of study patients for inclusion

**Table 1:** Demographic details of included participants.

|  | **Survivors**<br>**n=15** | **Non-survivors**<br>**n=13** | **Univariate odds ratio**<br>**(95%CI)**<br>***p*-value** |
| --- | --- | --- | --- |
| **Median age in years (IQR)** | 32 (26-44) | 36 (26-46) | 1.0 (0.96 : 1.07) 0.75 |
| **Male Gender (n, %)** | 7 (46%) | 9 (69%) | 2.5 (0.54 : 12.1) 0.23 |
| **HIV co-infection (n, %)** | 12 (86%) | 10 (76%) | 0.25 (0.02 – 2.8) 0.26 |
| **Estimated BMI** | 22.4 (20.7-23.2) | 22.4 (19-23.5) | 0.90 (0.68 : 1.1) 0.44 |
| **Median CD4 count (cells/mm<sup>3</sup>) (IQR)** | 128 (25-290) | 255 (n=1) | 1.0 (0.98 : 1.04) |
| **Median Glasgow Coma Score (IQR)** | 14 (13-15) | 11 (6-13) | 0.13 (0.22 : 0.8) 0.02 |
| **Median CSF WCC (cells/mm<sup>3</sup>) (IQR)** | 28 (0-685) | 33 (7-209)<br>n=8* | 0.95 (0.4 : 2.2) 0.91 |
| **Median CSF bacterial load (DNA copies/ml) (IQR)** | 5.7E+05 (3206 – 4.7E+06) | 4.37E+07<br>(8.43 E+06 – 2.18E+08) | 6.0 (1.2 : 29.2) 0.02 |
| **Median Duration of symptoms at presentation in hrs (IQR)** | 48 (24-72) | 48 (21 – 60) | 0.99 (0.68 : 1.02)<br>p= 0.66 |
| **Death day 10 post admission** | 0/15 (0%) | 10/13 (76%) | ref |

**Table 2:**
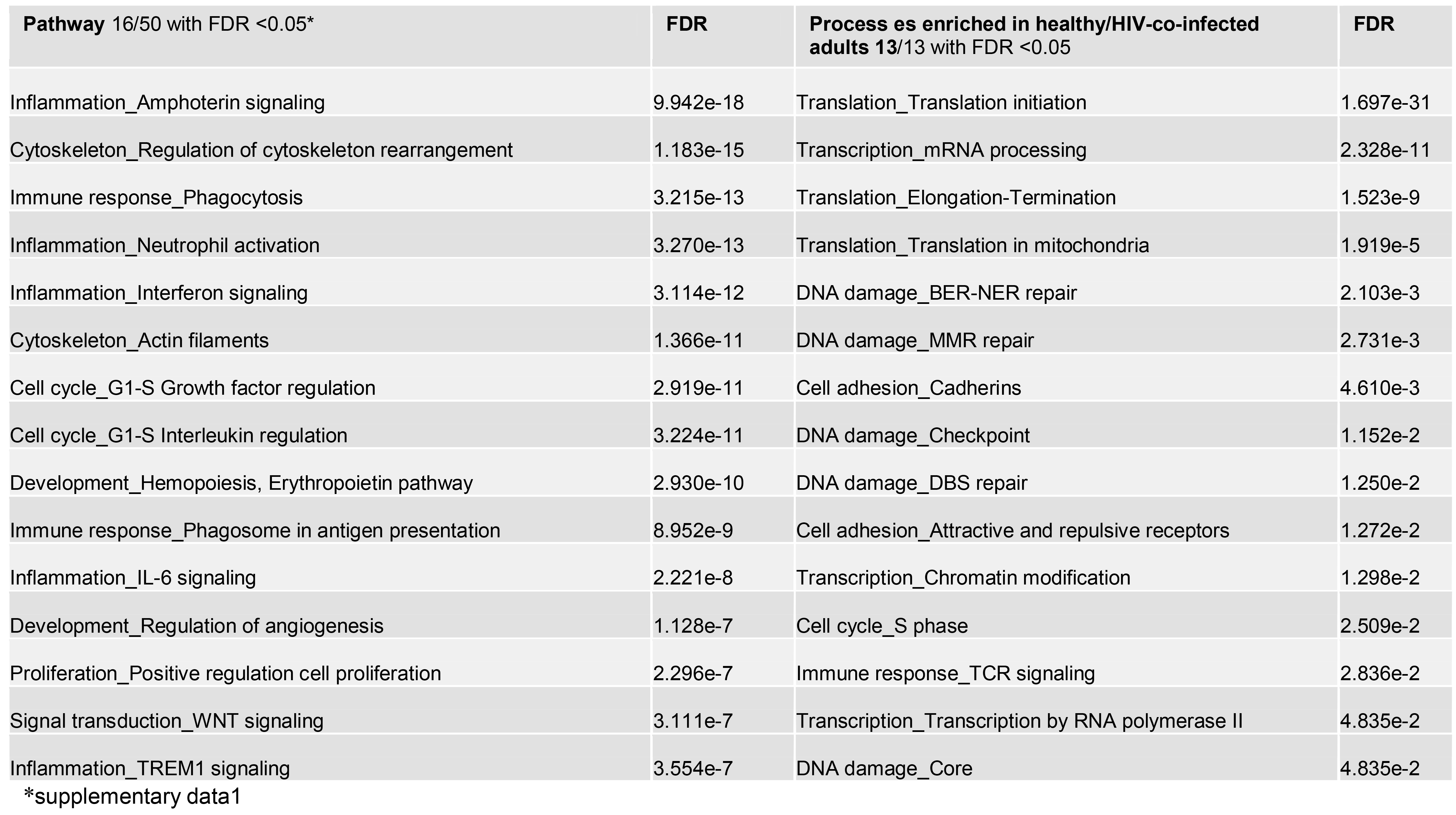
METACORE pathways & processes enriched in blood compartment of patients with PM compared to healthy/HIV-coinfected adults.

### CSF in PM is highly enriched for both PMNs and mast cells

To investigate the relatively lower reported CSF WCC in the patients with PM in this study compared to patients in higher income settings^32^ (Table 1), we first inferred the cellular composition of CSF in detail using CIBERSORTx^33^. Across samples, between 41-91% of cells were determined to be neutrophils (Figure 2). As an additional control for the CIBERSORTx analysis, we included two available transcriptomes from *S. pneumoniae* cultured *in vitro* CSF in the presence of freshly purified neutrophils in the analysis^34^, finding that CIBERSORTx correctly identified these transcriptomes as primary neutrophils. Other cell types detected in the PM-CSF samples included an unexpectedly high proportion of activated mast cells (up to 40% of cells / sample). Other cell types comprised 5-10% of the samples, but the subsets found were highly heterogenous (Figure 2).

**Figure 2:**
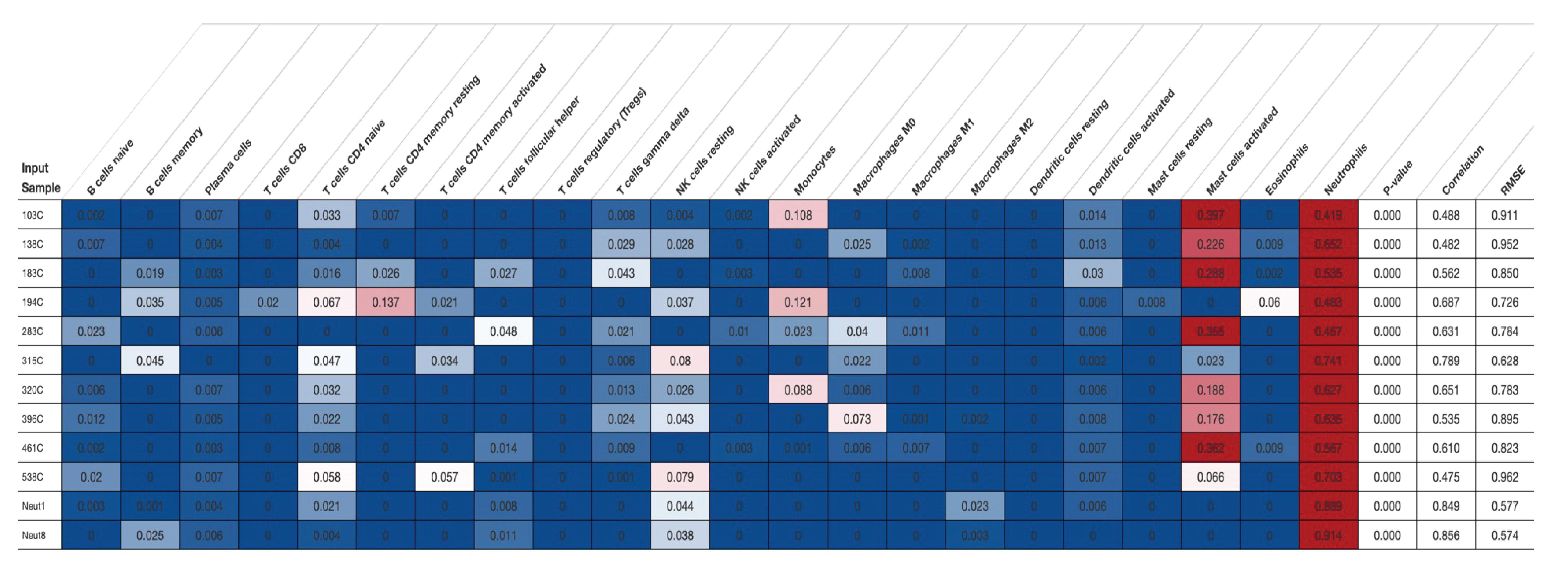
CSF composition is enriched for granulocyte populations of neutrophils and activated mast cells in pneumococcal meningitis. Cellular deconvolution of the CSF transcriptome in patients with PM using CIBERSORTx. Rows represent individual CSF samples from a unique patient. Neut1 and Neut2 indicate transcriptomes from purified human neutrophils incubated in normal CSF. Columns indicate cell-type. Numbers (and associated colours) show enrichment percentage of gene expression unique to that cell type. Adjusted p-value and correlation matrix denoted per sample.

### The inflammatory response is primarily compartmentalised within the CSF in PM

To investigate the degree by which the inflammatory response localises to the CNS or represents a systemic response to infection in PM, we compared differential gene expression between paired patient blood and CSF. The transcriptomes for the two compartments clustered separately on PCA (Figure 3A), with 2293 genes significantly differentially expressed in the CSF compartment (LFC >2, FDR <0.05) compared to 909 in the blood compartment (Figure 3B). Heatmaps of differential gene expression demonstrated variable expression of individual genes across the CSF samples (Figure 3C). Gene-level functional annotation using gene set enrichment analysis (GSEA) was used to determine if particular gene sets associated with functional processes (Gene Ontology, GO terms) were enriched in the different compartments. We found NfKb signalling (8 gene sets), CXCL8 and ICAM-1 activation (4 gene sets each) were enriched in the CSF compartment, whereas in blood we found evidence of enrichment of TUBA4A microtubular formation and MAP2 Kinase signalling (6 gene sets) (Figure 3D). We then tested for enrichment of particular REACTOME pathways within the GSEA analysis, finding upregulation of pathways involving TLR-9, MyD88 and cytokine activation in the CSF, and pathways involving fibrin activation, haem scavenging and metabolic activity in the blood (Figure 3E). To further characterise the transcriptional activity in both compartments, we undertook an upstream analysis using MetaCore (Clarivate analytics). Of 100 pathways/processes identified, >75% of upstream processes in CSF involved upregulation of genes involved in the innate response, including inflammasome activation, cytokine signalling, neutrophil activation, phagocytosis and leukocyte chemotaxis (Table 3A, supplementary Figure 1A). In contrast in blood pathways analysis identified processes involved in platelet activation, NETosis, cell cycle, cell adhesion, IL-6, IL-4 activation and antigen presentation activity (Table 3B, supplementary Figure 1B).

**Figure 3:**
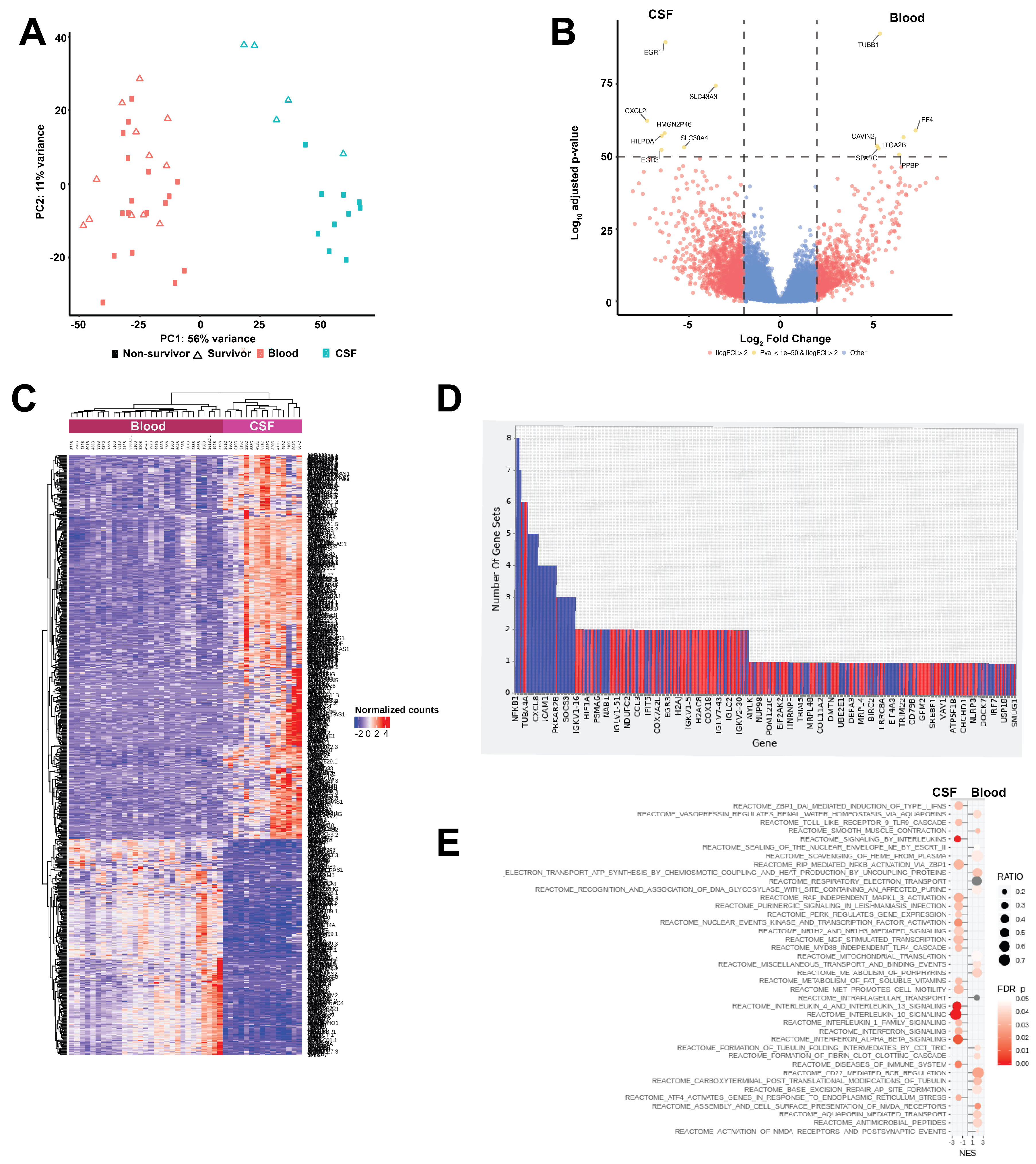
The inflammatory response is primarily compartmentalised within the CSF in adults with PM. **(A) :** Principal component analysis of gene expression between CSF (red) and blood (blue) transcriptomes in patients with PM. **(B):** Differential gene expression between the blood and CSF compartments of patients with PM. Yellow dots indicated genes above the significance threshold (FDR <1x1050, y axis, Log2 fold change (LFC) >2, x axis), red dots indicate differentially expressed genes above the fold change threshold (LFC >2.0), but below FDR <1x1050). **(C):** Heatmap showing heterogeneity of differential gene expression across samples between blood (red) and CSF (purple). Rows show individual genes, columns are individual patients, expression by normalized counts in transcripts per million (TPM) denoted by scaled colour bar. **(D):** Enrichment of individual genes within gene ontology (GO) sets between CSF (blue) and blood (Red) using GSEA. **(E):** Enrichment of pro-inflammatory pathways in the CSF of patients with PM (right) compared paired blood (left) using gene-set enrichment analysis (GSEA) and REACTOME, stratified by Normalised enrichment score (NES). Dot size indicates the ratio of upregulated genes within each pathway, colour denotes false discovery rate (FDR)/adjusted p value above the significance threshold.

**Table 3A:** METACORE pathways & processes enriched in CSF compartment of patients with PM compared to blood.

| **Pathway** 13/100 with FDR <0.05 | **FDR** | **Process** 13/93 with FDR <0.05 | **FDR** |
| --- | --- | --- | --- |
| Immune response_IL-1 signaling pathway | 1.662e-16 | Inflammation_Amphoterin signaling | 1.206e-15 |
| Immune response_IFN-alpha/beta signaling via JAK/STAT | 1.662e-16 | Immune response_Phagocytosis | 2.976e-15 |
| Immune response_HSP60 and HSP70/ TLR signaling pathway | 1.402e-15 | Immune response_Phagosome in antigen presentation | 6.469e-14 |
| Immune response_CD40 signaling in dendritic cells, monocytes, and macrophages | 1.505e-14 | Inflammation_Neutrophil activation | 2.237e-13 |
| Immune response_IL-18 signaling | 1.505e-14 | Inflammation_Protein C signaling | 5.954e-11 |
| Oxidative stress_ROS-induced cellular signaling | 1.505e-14 | Inflammation_IL-10 anti-inflammatory response | 5.954e-11 |
| Immune response_Plasmin signaling | 2.330e-14 | Inflammation_MIF signaling | 1.096e-10 |
| Immune response_IL-3 signaling via JAK/STAT, p38, JNK and NF-kB | 3.141e-14 | Inflammation_Interferon signaling | 1.096e-10 |
| Immune response_C5a signaling | 2.138e-13 | Cell cycle_G1-S Interleukin regulation | 1.721e-10 |
| Signal transduction_CXCR4 signaling via MAPKs cascades | 2.278e-13 | Development_Regulation of angiogenesis | 1.004e-9 |
| TNF-alpha and IL-1 beta-mediated regulation of contraction and secretion of inflammatory factors | 2.489e-13 | Development_Hemopoiesis, Erythropoietin pathway | 1.182e-9 |
| Neurogenesis_NGF/ TrkA MAPK-mediated signaling | 2.899e-13 | Inflammation_TREM1 signaling | 1.182e-9 |
| Signal transduction_NF-kB activation pathways | 3.425e-13 | Inflammation_IL-12,15,18 signaling | 3.321e-9 |

**Table 3B:**
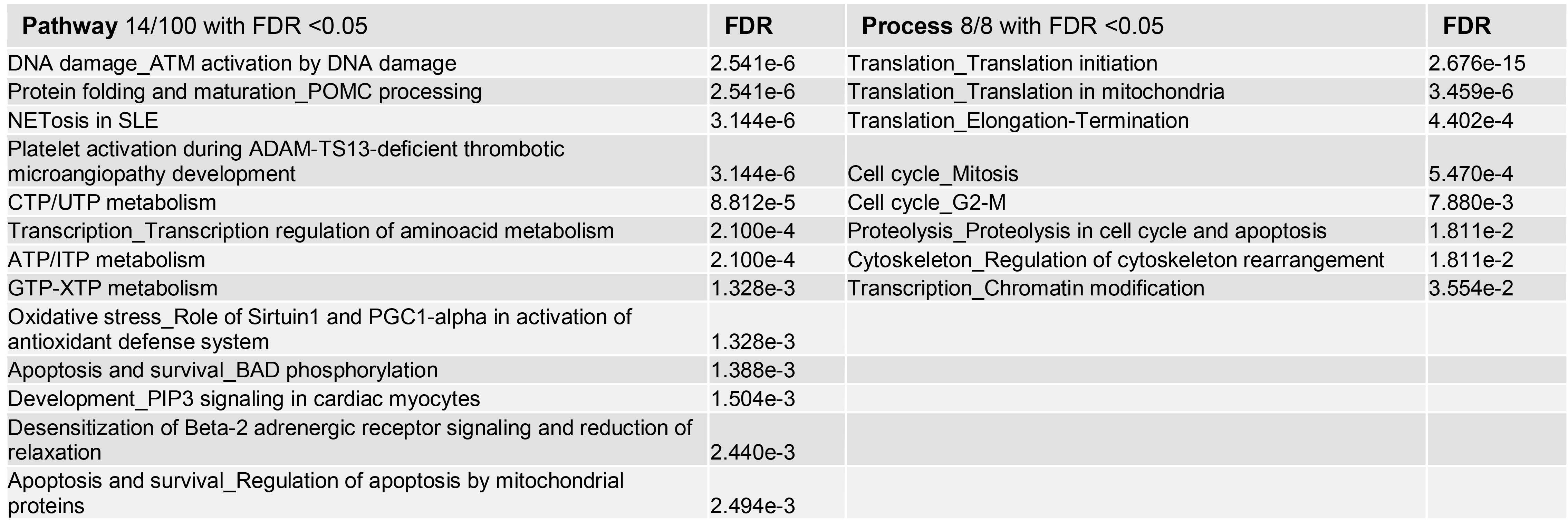
METACORE pathways & processes enriched in blood compartment of patients with PM compared to CSF.

### CSF transcriptomes in PM patients show significant differences between survivors and non-survivors

To further interrogate the inflammatory response and investigate processes associated with poor outcome, we tested for evidence of differential gene expression between survivors and non-survivors in both the CSF and blood compartments using DESeq2. After ruling out the existence of batch effects, we found separation of CSF transcriptomes between survivors (n=5) and non-survivors (n=8) (Figures 3A, 4B). In CSF 1829 genes were differentially expressed (FDR <0.05 LFC >2) between survivors and non-survivors, with 1085 genes with increased expression in non-survivors and 744 genes with increased expression in survivors (Figure 4C, Supplementary Data 1). Expression of the neutrophil persistence or survival-associated genes *CSF3* and *NR4A3* were markedly enriched in non-survivor CSF (LFC >2.5, FDR <0.05)^35,36^ with proteoglycan *DCN*, Endothelin 1 (*EDN1*), Anti-sense organic ion carrier protein *SLCO4A1-AS1*, G0/G1 switch regulatory protein *FOSB*, Hypoxia-inducable lipid protein HILPDA, and prostaglandin endoperoxidase Synthase *PTGS2* (Figure 4C, Supplementary Data 1) providing evidence of hypoxic endothelial inflammation and proteolysis in the CSF of non-survivors. In survivor CSF, only the transcription factor *MYCL* was over-expressed to this threshold. In contrast, blood transcriptomes showed minimal differential gene expression between survivors or non-survivors (Figure 3A), with only 135 genes met the lower statistical threshold (FDR <0.05) for differential expression between outcome groups (52 genes upregulated in survivor blood and 83 in non-survivor blood).

**Figure 4:**
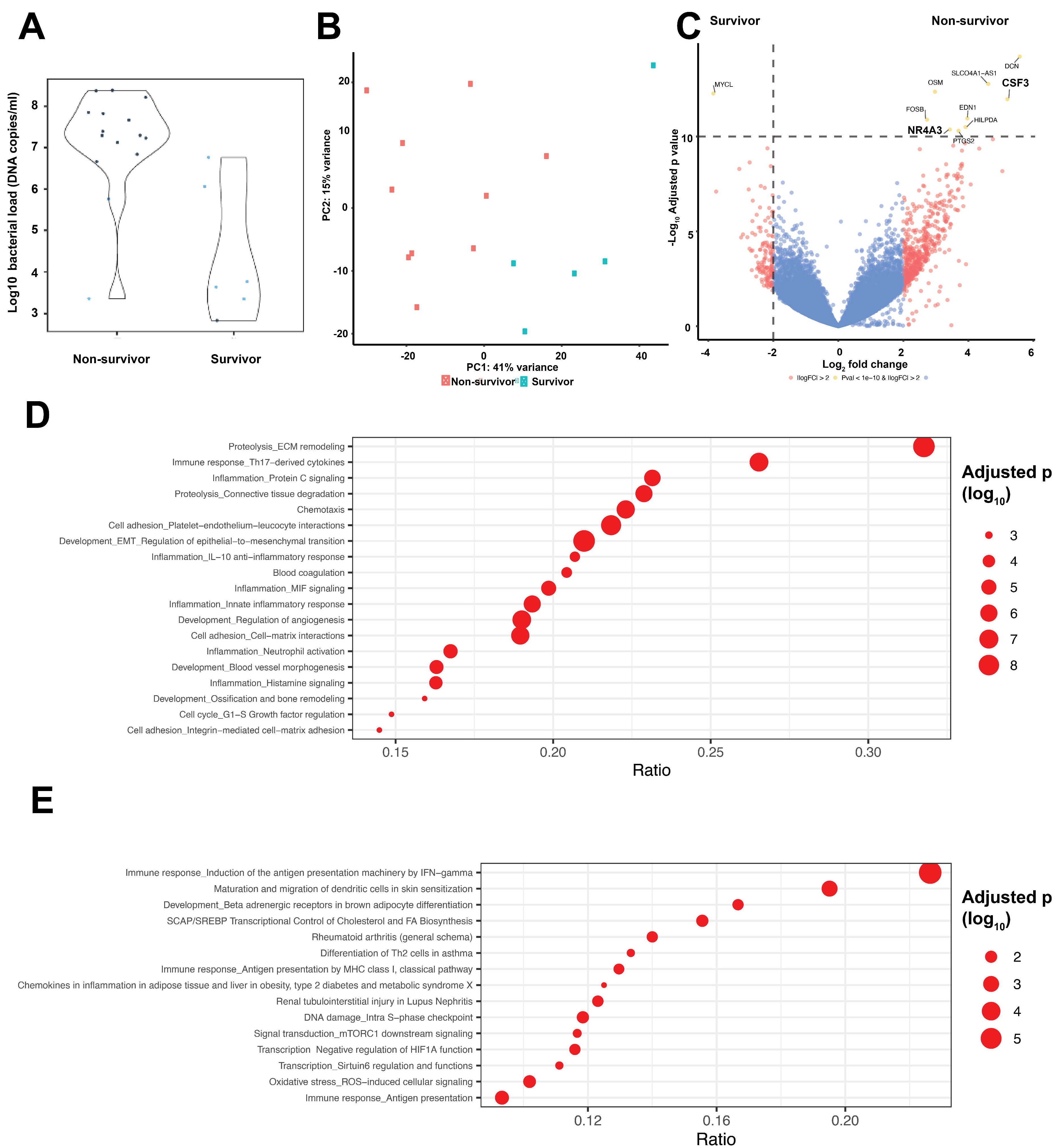
CSF in non-survivors of PM is enriched for IL-17, proteolysis, endothelial activation and innate inflammatory mediators. **(A):** CSF bacterial load (log_10_ DNA copies/ml, y axis) between non survivors and survivors of PM. **(B)**: Extent of differences in the CSF transcriptome between non-survivors (red) and survivors (green) on PCA **(C)**: Differential gene expression between the CSF of survivors and non-survivors of PM. Yellow dots indicated genes above the significance threshold (FDR p<1x1010, y axis, Log2 fold change (LFC) >2, x axis), red dots indicate differentially expressed genes above the fold change threshold (LFC >2.0), but below FDR <1x1010). **(D):** Upstream pathway analysis of the CSF transcriptome in non-survivors of PM using METACORE. Dot size indicates the adjusted p value/FDR below the significance threshold (<0.05), ratio indicates the proportion of genes in each process enriched in the differentially expressed genes **(E):** Upstream pathways analysis of the CSF transcriptome in survivors of PM using METACORE.

Using GSEA and REACTOME, we identified multiple pathways that were highly enriched (FDR <0.05) in non-survivor CSF, including those involved in smooth muscle contraction, activation of matrix metalloproteinases, phagocytosis, CD22 dependent BCR signalling, and neutrophil degranulation (Supplementary Figure 1C). In contrast, CSF transcriptomes from survivors were enriched for PD-1 signalling and second messenger molecules. We tested for interactions between individual highly expressed genes in survivor and non-survivor CSF using a network analysis in XGR, visualised using Gephi [http://gephi.org]. We found a small network of co-expressed genes in survivor CSF that included *HIST2H4A*, *CCND1* and *PIK3CD*, suggesting DNA and cellular damage response and immunological receptor signalling (Supplementary Figure 3A). In contrast, the non-survivor CSF gene network was dominated by a central cluster of pro-inflammatory cytokine genes including *TNF, IL1-b, MMP9* and *IL6*, with adjacent clusters of genes coding for vasoconstriction, platelet aggregation and cytoskeleton remodelling (Supplementary Figure 3B).

Finally, we used MetaCore (Clarivate Analytics) to describe upstream processes that associate with the inflammatory responses observed between two outcome groups in CSF. We identified 16 enriched primary processes in the CSF of non-survivors, the majority related to proteolysis, Th-17 derived cytokines, platelet-endothelium-leukocyte interactions, connective tissue degradation, angiogenesis, and protein C and MIF signalling, suggesting profound cellular damage (Figure 4D, Table 4B). Contrastingly, the CSF of survivors was enriched for antigen presentation, ROS-induced signalling, HIF1A functions, antigen presentation and TH2 cell differentiation (Figure 4E, Table 4C).

**Table 4A:** Pathways identified by METACORE in blood gene expression in non-survivors.

| Pathway | FDR | Ratio |
| --- | --- | --- |
| IgE-dependent production of pro-inflammatory mediators by neutrophils in asthma | 4.128e-4 | 5/38 |
| Neutrophil-derived granule proteins and cytokines in asthma | 7.540e-4 | 5/49 |
| Eosinophil granule protein release in asthma | 8.190e-4 | 5/54 |
| Development_G-CSF-induced myeloid differentiation | 1.334e-3 | 4/30 |
| Development_Transcription regulation of granulocyte development | 1.390e-3 | 4/32 |
| NF-kB-, AP-1- and MAPKs-mediated proinflammatory cytokine production by eosinophils in asthma | 3.827e-3 | 4/43 |
| NETosis in SLE | 2.821e-2 | 3/31 |
| Immune response_IL-11 signaling via JAK/STAT | 3.254e-2 | 3/34 |

**Table 4B:** Pathways & processes identified by METACORE in CSF gene expression in survivors.

| Pathway (15/15) | FDR | Ratio |
| --- | --- | --- |
| Immune response_Induction of the antigen presentation machinery by IFN-gamma | 1.422e-6 | 12/53 |
| Maturation and migration of dendritic cells in skin sensitization | 9.450e-4 | 8/41 |
| Oxidative stress_ROS-induced cellular signaling | 7.155e-3 | 11/108 |
| DNA damage_Intra S-phase checkpoint | 9.381e-3 | 9/76 |
| SCAP/SREBP Transcriptional Control of Cholesterol and FA Biosynthesis | 9.482e-3 | 7/45 |
| Renal tubulointerstitial injury in Lupus Nephritis | 1.348e-2 | 8/65 |
| Rheumatoid arthritis (general schema) | 1.424e-2 | 7/50 |
| Development_Beta adrenergic receptors in brown adipocyte differentiation | 1.448e-2 | 6/36 |
| Transcription_Negative regulation of HIF1A function | 1.448e-2 | 8/69 |
| Immune response_Antigen presentation by MHC class I, classical pathway | 1.702e-2 | 7/54 |
| Signal transduction_mTORC1 downstream signaling | 3.037e-2 | 7/60 |
| Differentiation of Th2 cells in asthma | 3.523e-2 | 6/45 |
| Transcription_Sirtuin6 regulation and functions | 3.523e-2 | 7/63 |
| Chemokines in inflammation in adipose tissue and liver in obesity, type 2 diabetes and metabolic syndrome X | 4.431e-2 | 6/48 |
| G-protein signaling_N-RAS regulation pathway | 4.836e-2 | 5/33 |
| PROCESS: Immune response_Antigen presentation | 3.683e-3 | 18/193 |

**Table 4C:** Pathways and processes identified by METACORE in CSF gene expression in non-survivors.

| Pathway | FDR | Ratio | Process | FDR | Ratio |
| --- | --- | --- | --- | --- | --- |
| IL-1 beta- and Endothelin-1-induced fibroblast/myofibroblast migration and extracellular matrix production in asthmatic airways | 4.281e-15 | 21/40 | Development_EMT_Regulation of epithelial-to-mesenchymal transition | 1.697e-9 | 47/224 |
| Cell adhesion_ECM remodeling | 3.384e-14 | 23/55 | Proteolysis_ECM remodeling | 1.697e-9 | 27/85 |
| Transcription_HIF-1 targets | 4.263e-14 | 29/95 | Inflammation_Amphoterin signaling | 1.748e-9 | 32/118 |
| IL-1 signaling in melanoma | 1.565e-12 | 19/42 | Cell adhesion_Platelet-endothelium-leucocyte interactions | 2.018e-8 | 38/174 |
| Immune response_IL-1 signaling pathway | 3.611e-12 | 25/82 | Immune response_Th17-derived cytokines | 1.275e-7 | 26/98 |
| Role of fibroblasts in the sensitization phase of allergic contact dermatitis | 8.123e-12 | 14/22 | Development_Regulation of angiogenesis | 1.459e-7 | 42/221 |
| Inflammatory mechanisms of pancreatic cancerogenesis | 1.820e-11 | 22/67 | Chemotaxis | 2.765e-7 | 31/139 |
| Th2 cytokine- and TNF-alpha-induced profibrotic response in asthmatic airway fibroblasts/ myofibroblasts | 8.700e-11 | 19/52 | Cell adhesion_Cell-matrix interactions | 2.765e-7 | 40/211 |
| Role of stellate cells in progression of pancreatic cancer | 1.252e-10 | 20/60 | Proteolysis_Connective tissue degradation | 1.094e-6 | 27/118 |
| Immune response_IL-17 signaling pathways | 1.252e-10 | 20/60 | Inflammation_Innate inflammatory response | 1.094e-6 | 35/181 |
| Development_Regulation of epithelial-to-mesenchymal transition (EMT) | 4.485e-10 | 20/64 | Inflammation_Protein C signaling | 2.190e-6 | 25/108 |
| Signal transduction_PDGF signaling via MAPK cascades | 5.688e-10 | 20/65 | Inflammation_MIF signaling | 1.090e-5 | 28/141 |
| TGF-beta-induced fibroblast/ myofibroblast migration and extracellular matrix production in asthmatic airways | 1.017e-9 | 19/60 | Inflammation_Neutrophil activation | 2.184e-5 | 36/215 |
| Immune response_Histamine H1 receptor signaling in immune response | 1.031e-9 | 17/47 | Development_Blood vessel morphogenesis | 2.829e-5 | 37/227 |
| Immune response_HSP60 and HSP70/ TLR signaling pathway | 1.095e-9 | 18/54 | Inflammation_Histamine signaling | 5.107e-5 | 35/215 |
| Immune response_CCL2 signaling | 1.095e-9 | 18/54 | Blood coagulation | 2.989e-4 | 19/93 |

Taken together, we show that the the inflammatory response in PM differs between blood and CSF in a cohort of predominately HIV-infected Malawian adults, with marked differences in the pre-antibiotic compartmentalised CSF inflammatory responses in patients who subsequently survived and those who died.

## Discussion

Adjunctive treatment with broadly effective anti-inflammatory agents such as dexamethasone has failed to improve the poor outcome of PM in LMICs^19,37^ and research into newer treatments has progressed slowly^38,39^. Here, we provide the first description of the paired CSF and blood transcriptome responses in patients with *S. pneumoniae* meningitis and identified host responses associated with lethal infection. We found the compartmentalised CSF response in non-survivors showed marked differences in gene expression to survivors, with enhanced activity of pathays involved in remodelling of the extracellular matrix, IL-17 responses, and vascular inflammation including platelet adhesion and activated protein C.

Many of the upregulated gene transcripts in the CSF of non-survivors correspond to CSF proteins associated with poor outcome from meningitis in previous studies, including tissue collagenases and matrix-metalloproteinases MMP 8 & 9, helping to validate our transcriptional findings^40–45^. Both IL-17 and T1-IFN driven mechanisms are important components of the mucosal host response to *S. pneumoniae* and may be associated with neutrophil recruitment across the blood brain barrier^46–50^. For example, IL-17 induces MMP-8 & 9 and vaso-active mediators (e.g. VEGF) at the mucosal surface to facilitate neutrophil mediated bacterial clearance^44,51,52^. In the CNS, IL-17 contributes to rapid BBB damage via these same pathways^53,54^. While IL-17 activity has not previously been described in pneumococcal meningitis, our finding of upstream IL-17 activity in CSF in PM concurs with prior evidence of IL-17 activity in other neurological infections^55,56^. The source of IL-17 in the CNS across neuro-inflammatory conditions is not yet well described.

PM is an acute disease, with rapid clinical onset, and the CSF classically contains over 50% polymorphonuclear cells (usually nearer 90%) which are commonly assumed to be neutrophils.^57^ As expected, the RNAseq data indicated the CSF in PM was enriched with granulocytes, but unexpectedly mast cells were the second commonest cell type identifed. The presence of activated mast-cells have been reported in CSF in TBM^58^ but not in acute bacterial meningitis, where their functional role in pathogenesis is not known. Resident mast cells within the CNS can be found both within the meninges and cerebral vasculature^59,60^, may produce GM-CSF as part of cerebral inflammation in MS^61^, whereas cerebral vasculature resident mast cells regulate BBB integrity after ischaemic insults^62^, releasing proteolytic enzymes in response to IL-17 following acute stroke^63–66^.

Cerebral ischaemia is a common complication of both PM and TBM^67–69^. While our finding of extremely high expression of *CSF-3* (GM-CSF) and activation of both IL-17 and proteolysis pathways in patients who died requires mechanistic validation, this observation suggests mast-cell mediated neuro-vascular inflammation and proteolytic effects on BBB integrity could contribute to disease pathogenesis in PM.

The host-pathogen interaction between *S. pneumoniae* and neutrophil phagocytosis is the core component of PM pathogenesis^13,70–75^. While neutrophils are required for bacterial clearance, neutrophil activity has well-recognised secondary harmful effects on the brain that are associated with worse outcomes in stroke, brain injury and encephalitis as well as meningitis^13,76–79^. Our transcriptome analysis allows neutrophil activity to be described in greater depth than neutrophil cell counts or measuring neutrophil-associated proteins in PM^23,80^. We found significantly increased expression of genes reflecting the potential presence of a pro-inflammatory, non-apototic subset of neutrophils in non-survivors, including *CSF3/NR4A3.* Expression of these genes may be triggered by cytokine release from necrotic neutrophils^36^ which are associated with more severe inflammation and tissue damage in other tissues^35,36,81^ through release of IL-17 and TNF alpha from surrounding cells^82,83^. Despite a near-identical time to presentation, we also found considerably higher bacterial DNA copy numbers in the CSF of non-survivors compared to survivors, indicating a possible association between poor control of bacterial replication and outcome^15,17^. While our earlier report of the proteome in these same patients revealed extensive neutrophil-derived proteins in CSF, the range of neutrophil functional states in meningitis have not been previously described. We hypothesise that neutrophil dysfunction could contribute to poor control of *S. pneumoniae* replication, and potentially lead to a detrimental positive feedback as higher bacterial numbers would increase the concentration of bacterial factors associated with poor outcomes such as pneumolysin^80,84^ that disrupt phagocyte function, leading to extensive tissue damage and death^12,71,73,80,85,86^. While requiring further investigation, our findings provide important points for future investigation into the inflammatory mechansisms associated with death in PM.

The inflammatory processes and very high bacterial loads in our patients are more marked than reports from patients with PM in higher income settings, and therefore may not be as amenable to modulation by systemic glucocorticoid therapies^87–89^, providing possible explainations to why dexamethasone was ineffective in this setting^19^. Alternative adjunctive therapeutic approaches that have been proposed include selective blockade of the IL-17 and T1-IFN pathways to limit some damaging inflammatory elements in PM, reduction in damaging effects of NETosis using treatment with DNAse, or anti-inflammatory antibiotics including daptomycin^13,90–92^. However, in the presence of the complex interacting processes demonstrated in our study, these interventions may have unpredictable effects on outcome^93,94^. Further mechanistic studies using appropriate disease models are required before further clinical trials of adjunctive agents in PM in our setting.

## Limitations

Our data are derived from a prospective observational cohort of participants recruited to a clinical trial^1^. Our analyses of the host inflammatory response using proteomics^80^ and transcriptomics were conducted as exploratory secondary outcomes to provide data to inform the trial, and thus we were unable to power our study design *a priori* to detect differences between outcome groups. Further, the patients in this study were predominately HIV-1-infected, comparisons by HIV serostatus were not possible due to small numbers of HIV-negative patients. We were unable to stratify our analysis by viral load or CD4 count as these data were not included in the original trial protocol, and were unable to directly control for HIV serostatus in the blood transcriptome data. However, the differential gene expression in blood from our patients are qualitatively distinct to published transcriptomic data for PLWHIV in South Africa with and without tuberculosis infection^95^, suggesting our findings relate to invasive pneumococcal infection as opposed to chronic inflammation triggered by untreated HIV-infection. We were unable to find reports of the CSF transcriptome in PLWHIV without concurrent infection to undertake the same analysis.

All HIV-1-infected patients were WHO clinical stage 3. The relatively small numbers of patients with high quality CSF RNAseq libraries limited our ability to stratify transcriptome signatures by either neutrophil count or bacterial load. We controlled as far as possible for potential confounders and technical differences inherent in these types of comparisons through careful selection of participants and lengthy QC and re-mapping of the data, but we cannot fully control for inter-individual differences between patients. Hence we have limited the data presented to replicable and biologically plausible findings. The CIBERSORTx model used to analyse the cellular composition of CSF is trained on immune and not brain cells, neurons or cell components of the BBB in CSF may have been missed using this approach.

Further, specific transcriptional modules for critical neutrophil functions, such as phagocytosis and trans-endothelial migration are currently lacking, and instead we used the gene response-module analysis approach using GO terms that have high sensitivity but untested specificity. Sufficient CSF was not available to validate our transcriptomic findings at the protein level; but our findings are supported by our previous proteomic analysis^80^ and other biomarker studies^44,45,52^. Pre-hospital delay may have influenced the differences in bacterial load and transcriptional responses seen between survivors and non-survivors, although estimates of pre-hospital disease onset times were not associated with outcome in either our study or in a larger Malawi meningitis database^1,6^.

## Conclusions

Using RNAseq we have shown that inflammation is strongly compartmetalised between the CSF and blood in patients at presentation with PM, and identified multiple genes for which high CSF levels of expression were associated with poor outcome. These data identify potential biomarkers for poor outcome, and provide new avenues for investigation into the role(s) of activated mast cells, neutrophil functional state and IL-17 in the pathogenesis of PM. The crucial question of how best to protect the brain from both damage caused by pneumococcal cytolysins and host-derived, neutrophil mediated proteolytic damage should be prioritised for future discovery.

## Methods

### Participants

Adults and adolescents presenting to Queen Elizabeth Central Hospital in Blantyre, Malawi with subsequently proven bacterial meningitis caused by *S. pneumoniae* between 2011- 2013 were included (Current Controlled Trials registration ISRCTN96218197)^1^. All CSF and blood samples were collected prior to administration of parenteral ceftriaxone 2g BD for 10 days^19,20^. Clinical data are from the first recording on admission to hospital, follow up was done to six weeks post-discharge^1^.

### Procedures

Routine CSF microscopy, cell count, and CSF culture was done at the Malawi-Liverpool-Wellcome Trust Clinical Research Programme laboratory in Blantyre, Malawi as previously described^1^. Culture negatives samples were screened using the multiplex real-time polymerase chain reaction for *S. pneumoniae*, *N. meningitidis* and *Haemophilus influenzae type b* (Hib) kit from Fast-Track Diagnostics (FTD Luxemburg) according to the manufacturer’s instructions, bacterial loads were estimated from Ct values. We collected 2.5 ml of CSF and whole blood for transcriptional profiling in blood PAX-gene® (Pre-AnalytiX, Qiagen, USA) tubes, incubated for 4 hours at room temperature, and stored at -80 degrees Celsius. In-hospital HIV testing was done on all patients by the clinical teams using point-of care Genie™ HIV1&2 test kits (BioRad, USA).

RNA was extracted from blood and CSF using the PAXgene® Blood miRNA kit (Pre- Analytix, Qiagen, USA) according to the manufacturer’s instructions, with an additional mechanical disruption step in the CSF samples to disrupt the pneumococcal cell wall at 6200 rpm for 45 seconds in the Precellys evolution tissue homogenizer (Bertin Instruments). The extracted RNA was quantified and RNA Integrity Number (RIN) scores calculated using RNA Tapestation 4200® (Agilent, USA) and Nanodrop® (Thermoscientific, USA). Extracted RNA samples underwent library preparation for polyA tailed mRNA with a RNA concentration of >1ng/1ul using with Kapa RNA hyperPrep kit (Roche), followed by 75 cycles of Next-generation sequencing with NextSeq® (Illumina, USA) using a single flow cell to further minimise batch effects by the University College London Genomics Unit.

Human CSF samples were a kind gift from Professor Diederik van de Beek at the Amsterdam Medical Centre, University of Amsterdam, The Netherlands. Surplus normal lumbar CSF was obtained from diagnostic lumbar punctures with consent, snap frozen, shipped at -80°C and thawed on ice to preserve active complement. Complement was depleted from CSF where required by heating to 65°C for ten minutes.

Fresh human neutrophils were extracted from whole blood of healthy lab donors by negative selection using the MACSxpress® system (Miltenyibiotec, USA) according to the manufacturer’s instructions. Erythrocytes were depleted post neutrophil isolation by incubation for 8 minutes with 1X Invitrogen RBC lysis buffer (ThermoFisher, USA) prior to all experiments. Neutrophil viability was assessed by Trypan blue staining. Neutrophils were counted using a cell chamber and adjusted in all experiments to 2x10^6^ cells/ml. Neutrophils were re-suspended in HBSS with 10% FBS and kept at 37°C until use (<4 hrs). Neutrophils were incubated in warmed CSF for 30 minutes in 5% CO^2^ at 37°C, before pelleted and re- suspended into 1 mls of RNA*later.* All samples were incubated for 4 hours at room temperature and then frozen at -80°C. RNA was extracted following a method developed by Mann et al, using the MirVana phenol based extraction kit (Thermofisher) as previous reported as part of a protocol to investigate the *in vitro* pneumococcal transcriptome.^34,95^ Briefly, samples were thawed, pelleted and RNA protection media was removed. Cell lysis buffer was applied, samples were placed in a FastPrep MatrixE tube, undergoing mechanical cell wall disruption in a Precellys machine speed of 6200 rpm for 45 seconds. Homogeneates were incubated in a water bath for 10 minutes at 70°C, cooled on ice and then pelleted at 12k x g, 5 min, 4°C in pre-cooled microfuge. The supernatant containing RNA was removed and passed through a Qiashredder (Qiagen) for 2 minutes at 12k x g, in the same 4°C in pre-cooled microfuge.

RNA extraction was completed using the MirVana kit (ThermoFisher) following the manufacturers instructions. Following nucleic acid extraction, TurboDNAase enzyme and buffer (1:10 ratio) (ThermoFisher) were added to each sample and incubated at 37°C for 30 minutes. RNA quality was quantified using the Tapestation/BioAnalyser. Ribosomal RNA was depleted using human rRNA depletion beads (NewEnglandBiolabs). Libraries for all samples were prepared simultaneously to minimise batch effects as previously^34^.

Ribodepletion was done with the Illumina RiboZero Gold kit, following the manufacturers instructions. Extracted RNA samples underwent library preparation for total RNA sequencing in samples where the pre-ribodepletion RNA was >1ng/1ul using with Kapa RNA hyperPrep kit (Roche), followed by 75 cycles of Next-generation sequencing with NextSeq® (Illumina, USA) by the Pathogen Genomics Laboratory at University College London.

### Bioinformatics and statistical analysis

All conventional statistical tests were two tailed, alpha <0.05 determined statistical significance. 95% confidence intervals are presented for odds ratios. Logistic regression was used to model associations between clinical outcomes and risk factors while controlling for confounding factors.

Sequence data quality was assessed prior to mapping by using the FASTQC toolkit (http://www.bioinformatics.babraham.ac.uk/projects/fastqc/). Mapping quality and percentage of properly mapped pairs, assessed from the BAM files, were considered for quality control. Sequenced cDNA libraries were mapped at the transcript and gene levels to the human genome (assembly GRCh38) using *Salmon* v0.8.2 (https://salmon.readthedocs.io/en/latest/salmon.html). We removed mapped genes for specific haemoglobin processes prior to data analysis^96^. ^96^. Detected haemoglobin levels were similar between biological groups, and did not constitute a confounding factor.

For all differential expression analyses, we first normalised and compared gene expression using the R package DESeq2 was used to test for differential gene expression on log_2_ normalised gene counts^97^. False Discovery Rate (FDR) corrected p-value <0.05 was used to threshold for significance in the differential gene expression analysis. Significantly enriched pathways were denoted by FDR corrected p-value <0.05.

We then used Gene Set Enrichment Analysis (GSEA) to rank differentially expressed genes against the molecular signatures database for gene-ontology terms to analyse function-specific gene expression for individual cell types (http://software.broadinstitute.org/gsea/msigdb/index.jsp), and reported both gene sets and associated pathways identified with those gene sets in REACTOME.

Metacore (Clarivate.com) was used to test for overenrichment of upstream pathways and processes using differentially expressed genes in patients with PM. Metacore employs a hyper-geometric model to determine the significance of enrichments Immune cell composition was estimated from the RNA-seq gene expression data using CIBERSORTx, a computational tool that infers cell type abundances from bulk tissue transcriptomes^33^. The CIBERSORTx analysis was run using the default LM22 signature matrix consisting of 547 genes that distinguish 22 human hematopoietic cell phenotypes, including seven T cell types, naive and memory B cells, plasma cells, NK cells, and myeloid subsets.

Finally, we explored differentially expressed genes in an unbiased manner by testing for clusters of co-expressed genes in different compartments using R package XGR (http://galahad.well.ox.ac.uk:3020/subneter/genes). We visualised these clusters using Gephi (https://gephi.org/), analysing graphs by network centrality.

## Data Sharing

Mapped, sequence files for all included patients are available on a consent-basis through the European Phenome-Genome Archive at the European Bioinformatics Institute (EBI) https://www.ebi.ac.uk/ega/studies/EGAS00001003355

## Ethics

All participants or nominated guardians gave written informed consent for inclusion. Ethical approval for the transcriptomics study was granted by both the College of Medicine Research and Ethics Committee (COMREC), University of Malawi, (P.01/10/980, January 2011), and the Liverpool School of Tropical Medicine Research Ethics Committee, UK (P10.70, November 2010) Committee, Liverpool, UK.

## Author contributions

Conception or design of the work: ECW, JAGA, CV, GP, MaN, JSB, RSH

Data collection: ECW, JAGA, VM, MuN, DGL

Data analysis and interpretation: ECW, JAGA, PC, CV, GP, MaN, JSB, DGL, RSH

Drafting the article: ECW, JAGA, MaN, RSH, JSB

Critical revision of the article: ECW, JAGA, GP, CV, VSM, BD, MuN, DGL, MaN, JSB, RSH

Final approval of the version to be published: ECW, JAGA, PC, GP, CV, VSM, BD, MuN, DGL, MaN, JSB, RSH

## Acknowledgements

The authors would like to thank the study patients and guardians, the Bundles for Adult Meningitis (BAM) research team, clinical and laboratory staff at the Queen Elizabeth Central Hospital and Malawi-Liverpool-Wellcome Trust Clinical Research Programme in Blantyre Malawi for support given during the study and Professor Mike Levin and Dr Victoria Wright of Imperial College UK for support at the early stages of the project. We would like to thank Professor Judith Breuer and the staff of UCL Genomics (formerly Pathogen Genomics Unit) at University College London for their assistance with library preparation and RNA sequencing. The authors acknowledge the use of the UCL Legion High Performance Computing Facility (Legion@UCL), and associated support services, in the completion of this work. We would like to thank Professors Diederik van de Beek and Matthias Brouwer with the support of their clinical research teams at the Amsterdam Medical Centre for the kind donation of human CSF for this study.

## Supplementary Material

**Supplementary** Figure 1**: The compartmentalised inflammatory response in CSF during PM is driven by inflammasome-mediated proteins and neutrophil activity, in contrast with blood where oxidative stress and platelet activity dominate.**

**(A) :** Upstream pathway analysis of differentially expressed genes in the CSF transcriptome in PM when compared to blood using METACORE. Dot size indicates the adjusted p value/FDR below the significance threshold (<0.05), ratio indicates the proportion of genes in each process enriched in the differentially expressed genes
**(B) :** Upstream pathways analysis of differentially expressed genes in the blood transcriptome of patients with PM using METACORE. **(C):** REACTOME analysis using GSEA comparing pathways enriched between survivors and non-survivors of PM. Dot size indicates the ratio of differentially expressed genes to total genes in each pathway. Colour indicates adjusted p-value (False Discovery Rate - FDR). NES - Normalised Enrichment Score.

Supplementary Figure 2: Network analysis of differentially expressed genes in the CSF of survivors and non survivors.

Network analysis of significantly differentially expressed genes (FDR p-adj <0.05, LFC >2.0) in the CSF of survivors (A) and non-survivors (B) of pneumococcal meningitis. Gene clustering generated in XGR, graphs synthesised with Gephi, analysed with Wifan-Hu network centrality. Each node represents an individual gene, node size represents network centrality and connectivity. Edge thickness represents the strength of the relationship between nodes. Colours represent clusters by function and connectivity.

## Supplementary data

**Data 1:** Metacore results on pathways and processes enriched in blood vs CSF. **Data 2:** METACORE results pathways and processes enriched in survivor’s vs non- survivors

