## Supplementary Figure 2 for "The CSF transcriptome in pneumococcal meningitis reveals compartmentalised host inflammatory responses associated with mortality"

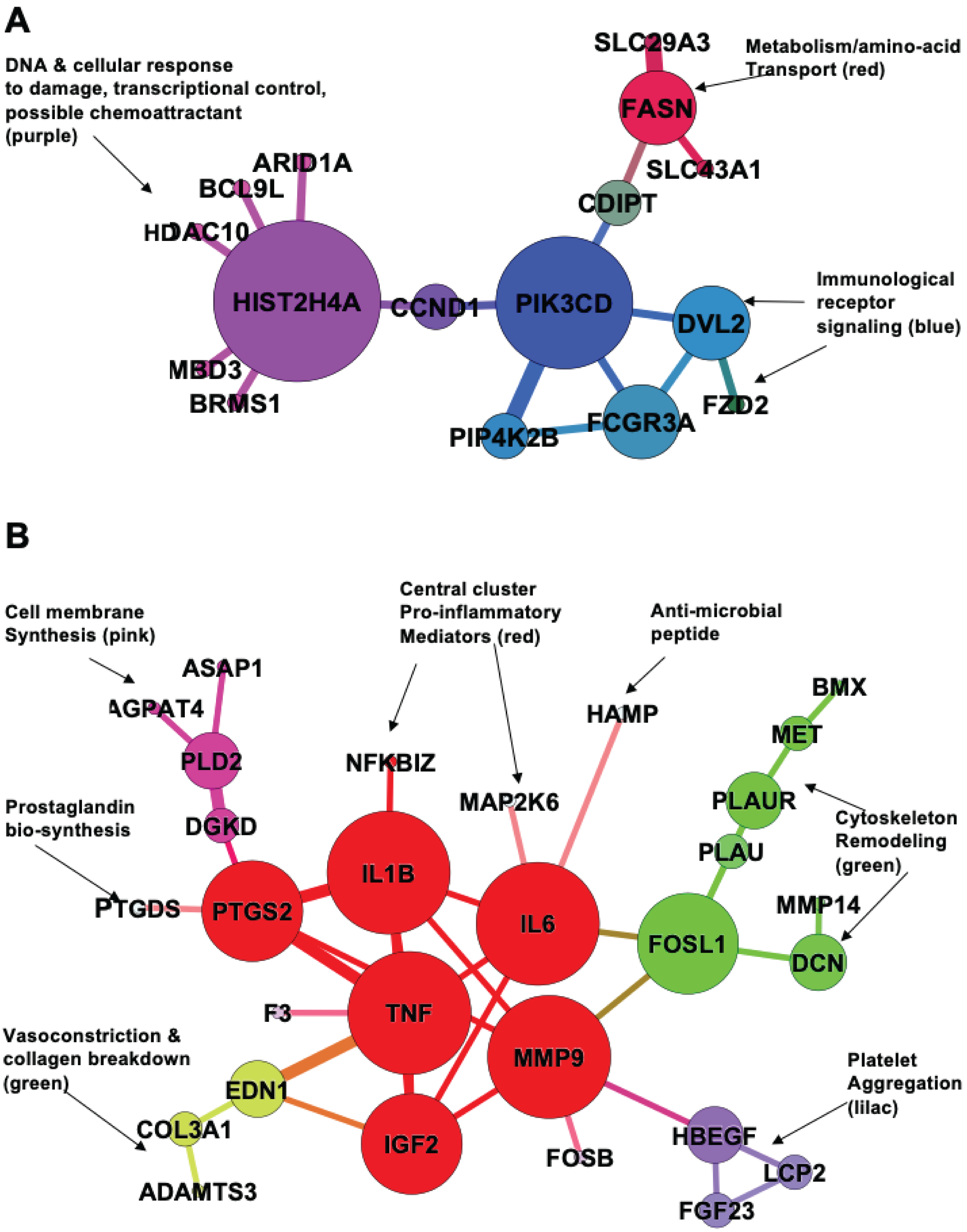


**Supplementary Figure 2: Network analysis of differentially expressed genes in the CSF of survivors and non survivors.**

Network analysis of significantly differentially expressed genes (FDR p-adj <0.05, LFC >2.0) in the CSF of survivors (**A**) and non-survivors (**B**) of pneumococcal meningitis. Gene clustering generated in XGR, graphs synthesised with Gephi, analysed with Wifan-Hu network centrality. Each node represents an individual gene, node size represents network centrality and connectivity. Edge thickness represents the strenth of the relationship between nodes. Colours represent clusters by function and connectivity.
