## Supplementary Figure 1 for "The CSF transcriptome in pneumococcal meningitis reveals compartmentalised host inflammatory responses associated with mortality"

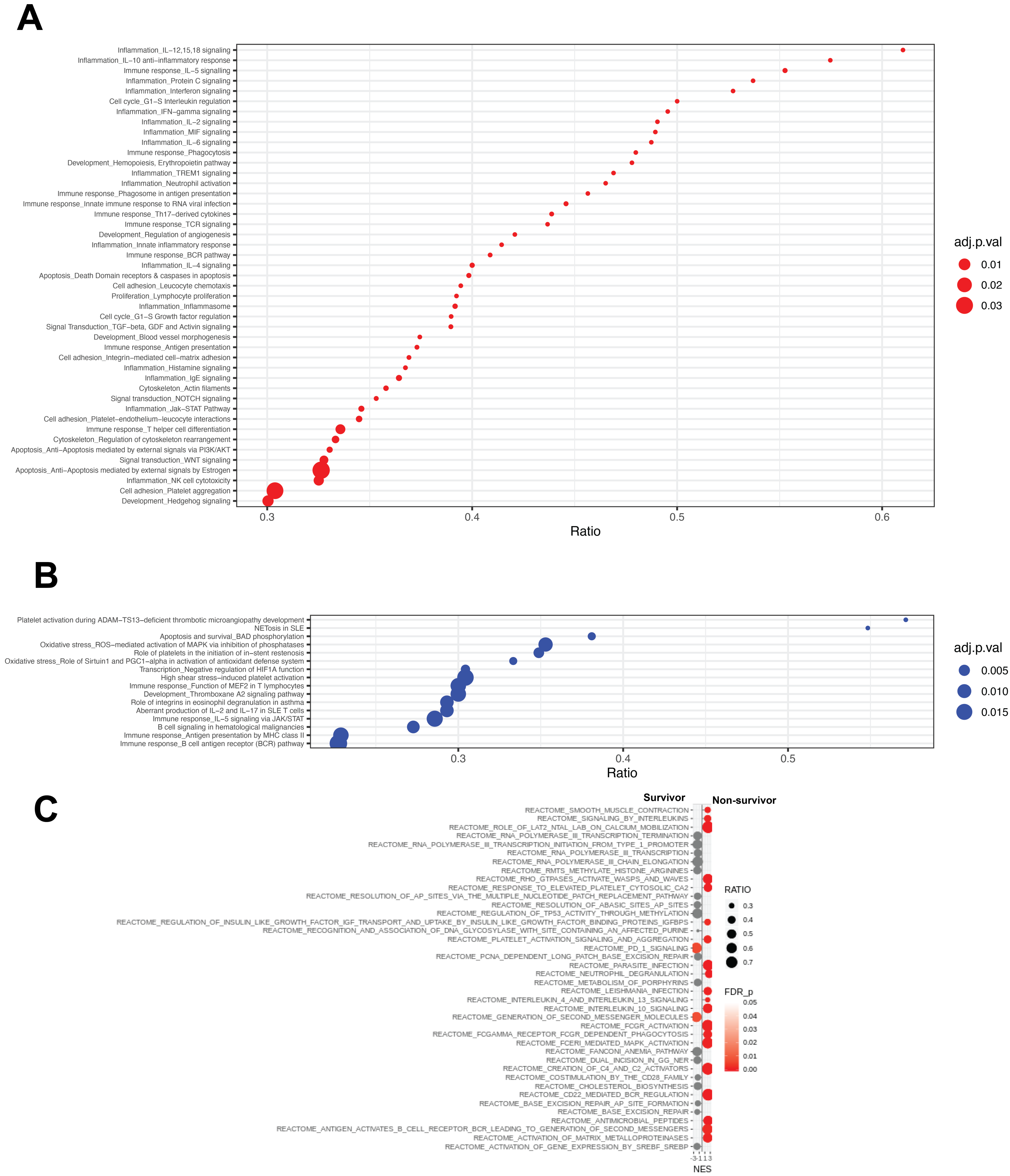


**Supplementary Figure 1: The compartmentalised inflammatory response in CSF during PM is driven by inflammasome-mediated proteins and neutrophil activity, in contrast with blood where oxidative stress and platelet activity dominate.**

**(A)**: Upstream pathway analysis of differentially expressed genes in the CSF transcriptome in PM when compared to blood using METACORE. Dot size indicates the adjusted p value/FDR below the significance threshold (<0.05), ratio indicates the proportion of genes in each process enriched in the differentially expressed genes **(B)**: Upstream pathways analysis of differentially expressed genes in the blood transcriptome of patients with PM using METACORE. **(C)**: REACTOME analysis using GSEA comparing pathways enriched between survivors and non-survivors of PM. Dot size indicates the ratio of differentially expressed genes to total genes in each pathway. Colour indicates adjusted p-value (False Discovery Rate - FDR). NES - Normalised Enrichment Score.
